# Savant: A Fast, Structure-Free Binding Affinity Predictor For Protein-Ligand Drug Screening

**DOI:** 10.64898/2026.11.15.766769

**Authors:** Sayal Guirales-Medrano, Colby T. Ford, Daniel Janies

## Abstract

Protein –ligand binding affinity determines which of many candidate molecules is worth synthesizing and assaying, making it the central measurement in early drug discovery. Binding affinity is commonly expressed as the dissociation constant, an equilibrium measure that indicates how easily a complex separates into its components. AI-based generative chemistry and make-on-demand chemical catalogs now produce molecules faster than rigorous method can evaluate them. Leading computational methods co-fold the complex before scoring the binding affinity between the protein and the ligand, which provides high accuracy but requires a graphics processing unit (GPU) and substantial time for every pair evaluated.

Here we present Savant, a machine learning model that estimates the dissociation constant using only the protein’s amino acid sequence and the ligand’s structure written as a line of text (SMILES). Savant needs no 3-dimensional folded structure, no docking, and no GPU. To test Savant fairly, we built a bench-mark designed to prevent any training data to leak into the testing set, drawing only on measurements published after the training data was assembled and on molecules chemically unlike anything the model had seen. On an independent benchmark, Savant ranks ahead of every other 3-dimensional structure-free method.

Savant is designed to rank compounds against a single protein, and performs as well as other methods that require a GPU, while running only on a central processing unit (CPU). Thus, Savant can screen libraries of 100,000 compounds in minutes with commodity hardware.

## Introduction

Bringing a new medicine to market is among the most expensive undertakings in applied science. Wouters et al. (1) estimated the median capitalized research and development investment per approved medicine at $985.3 million, with a mean of $1.3 billion, across 63 drugs approved between 2009 and 2018. Industry-sourced estimates are higher: reaching $2.5 billion capitalized per approval (2). Scannell et al. (3) documented that the number of new drugs approved per billion dollars of research and development spending has halved roughly every nine years since 1950. This trend represents an 80-fold decline in inflation-adjusted terms. Computational inputs have become dramatically cheaper over the same period while output has not improved. This relationship indicates a need for methods that triage drug candidates early and inexpensively (4).

That need has grown sharply because the supply of candidate molecules has outpaced our ability to evaluate them. Lyu et al. (5) docked 170 million make-on-demand compounds against two targets. Gorgulla et al. (6) created VirtualFlow which extended open-source screening past 1 billion compounds. The John Irwin lab created ZINC-22 which now catalogs tangible compounds at multi-billion scale (7). Artificial intelligence (AI) generative models add to the supply of protein–ligand structures, producing novel structures under property constraints (8, 9). This body of work results in a practical problem: 10^9^ to 10^10^ of available molecules but rigorous per-compound analyses take a long time to process. Sadybekov and Katritch (10) state that fast, cost-effective screening are the steps that will permit the evaluation of these libraries to approach the speed and scale at which they are being created.

If binding affinity between a protein and a small-molecule drug (ligand) can be predicted inexpensively, there is promise to resolve the triage problem. Affinity is commonly expressed through the dissociation constant (*K*_*d*_), which relates to the free energy of binding through Δ*G* = *−RT* ln *K*_*d*_ (11). Protein–ligand binding is governed by an interplay of hydrogen bonding, electrostatic interactions, hydrophobic effects, van der Waals interactions, and desolvation (11). There are practical limits of optimizing protein–ligand binding against molecular size motivated ligand efficiency as a design metric (12). Experimental determination of affinity of protein–ligand binding by surface plasmon resonance, isothermal titration calorimetry, or fluorescence-based assays remain the gold standards against which all computational work is calibrated. However, experimental assays require purified material and operate far below the throughput at which candidate molecules are now produced.

Physics-based computational methods span a wide accuracy and cost range, and neither end of that range solves the triage problem. At the inexpensive end, docking programs place ligands well but score them poorly. In a critical assessment of 10 docking programs and 37 scoring functions across 8 targets, no program or scoring function produced useful rank-order predictions of compound affinity (13). Moreover, the CASF-2016 comparative assessment shows that even best-case scoring correlations remain modestly successful (14).

At the expensive end, free-energy methods do achieve predictive accuracy, when tested across eight targets and 199 ligands (15). These methods perform well in prospective industrial projects (16). Free-energy perturbation, however, requires a high-quality 3-dimensional structure, a congeneric ligand series, and many GPU-hours per compound, which make this method intractable for library-scale triage.

Advances in computational 3-dimensional structure prediction promise to remove the first of those requirements. AlphaFold3 predicts biomolecular complexes including protein–ligand systems (17), and open co-folding models now predict 3-dimensional structure and binding affinity together (18–20). The evidence that this transfers to affinity is weaker than it first appears. Karelina et al. (21) found that although AlphaFold2 models capture binding-pocket structure well, ligand poses predicted from those models were not significantly more accurate than poses from traditional homology models. They were also substantially less accurate than poses predicted from experimentally determined structures. A predicted structure is therefore not yet a substitute for a measured one when the downstream task is quantitative.

A parallel line of work predicts affinity without any 3-dimensional structure at all. DeepDTA established that pKd can be predicted from sequence and SMILES alone (22). Subsequent methods refined the representation using molecular graphs (23), interaction transformers (24), contrastive protein-language embeddings (25), and bilinear attention with domain adaptation (26). PLAPT is the closest published peer to the present work, combining protein and chemical language-model embeddings in a small neural head (27). The featurizers themselves have improved independently: protein language models now recover structural information from sequence alone (28, 29). Chemical language models pretrained on tens of millions of SMILES provide transferable molecular representations (30, 31). Standardized collections support comparison across these methods (32).

How such methods are evaluated is itself contested, and this is central to the argument we make here. Chen et al. (33) showed that apparent deep-learning enrichment on the Directory of Useful Decoys-Enhanced (DUD-E) derived from analogue and decoy bias in the dataset rather than from learning protein–ligand interactions. Wallach and Heifets (34) demonstrated that most ligand-based classification benchmarks reward memorization rather than generalization. Sieg et al.

(35) formalized the bias controls needed to avoid the memorization problem when training machine learning algorithms. Volkov et al. (36) reached the harshest conclusion for affinity regression. Specifically, Volkov et al. find that an explicit description of protein–ligand noncovalent interactions provided no advantage over ligand-only or protein-only descriptors, and that memorization largely dominated true learning. Related analyses of graph networks reach compatible conclusions about what such embeddings encode (37). A single correlation pooled over many targets is vulnerable to this failure, because a model can score well by recognizing to which target a pair belongs. We separate the pooled correlation into a between-target component (many proteins to one ligand) and a within-target component (one protein to many ligands) and report them individually.

Any claim about predictive accuracy must also be read against the precision of the measurements. Kramer et al. (38) quantified the experimental uncertainty of heterogeneous public *K*_*i*_ data at a mean error of 0.44 and a standard deviation of 0.54 p*K*_*i*_. Kramer et al. also argued that this un-certainty defines a natural upper limit on achievable predictive performance, such that models reporting smaller errors are necessarily overtrained.

Here we present Savant, a gradient-boosted model that ranks ligands binding to a target protein through a predicted pKd from sequence and SMILES alone. Gradient-boosted trees were chosen because the feature set is heterogeneous tabular data combining embeddings, descriptors, and fingerprints, a regime in which tree ensembles remain competitive with deep networks and are robust to uninformative features (39–41). Our aim throughout was a model that runs on commodity hardware, which is the condition under which fast screening becomes broadly accessible (10).

## Methods

### Data acquisition and preparation

We sourced protein–ligand data from the public Hugging Face dataset jglaser/binding_affinity, compiled by Jens Glaser (42). The dataset aggregates records from BindingDB (43), PDBbind-CN (44, 45), BioLiP (46), and Binding MOAD (47). The dataset comprises 1.8 million pairs of protein amino acid sequences (`seq`) and ligand SMILES strings (`smiles`) with a measured affinity, provided both in micromolar units (`affinity_uM`) and as a negative logarithm in molar units (`neg_log10_affinity_M`).

We used pKd in molar units as the prediction target through-out, defined in Equation 1.

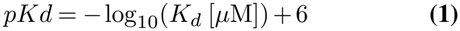

Two properties of this label govern the interpretation of every result reported below, and both are properties of the measurement rather than of any model.

First, the label pools assay types. The released file provides a single affinity column and no `standard_type` field, so dissociation constants (*K*_*d*_), inhibition constants (*K*_*i*_), and half-maximal inhibitory concentrations (IC_50_) are mixed and cannot be separated.

Second, the label is noisy, and we quantified the noise directly. We found that 11.9% of unique sequence and SMILES pairs appear more than once in the corpus, and that these replicates disagree by a pooled standard deviation of 0.81 pKd. A model that recovered the true value exactly would therefore still record a root mean squared error (RMSE) of approximately 0.81 against single measurements. This exceeds the approximately 0.5 p*K*_*i*_ uncertainty that Kramer et al. (38) reported for filtered public data. We read every error figure in this study against that floor, not against zero.

We collapsed replicate measurements of the same pair to their median before fitting, so that a frequently re-measured pair does not receive proportionally more weight than a singly measured one. We discarded pairs whose replicates disagreed by more than 1.5 pKd as irreconcilable. We also identified measurements recorded at the standard censoring concentrations of 10, 100, 1,000, and 10,000 *µ*M, which are reporting limits and not true measurements. With conversion to pKd we excluded measurements below 6 from the regression target while retaining them as negatives for classification, as seen in Fig. 1.

**Fig 1.**
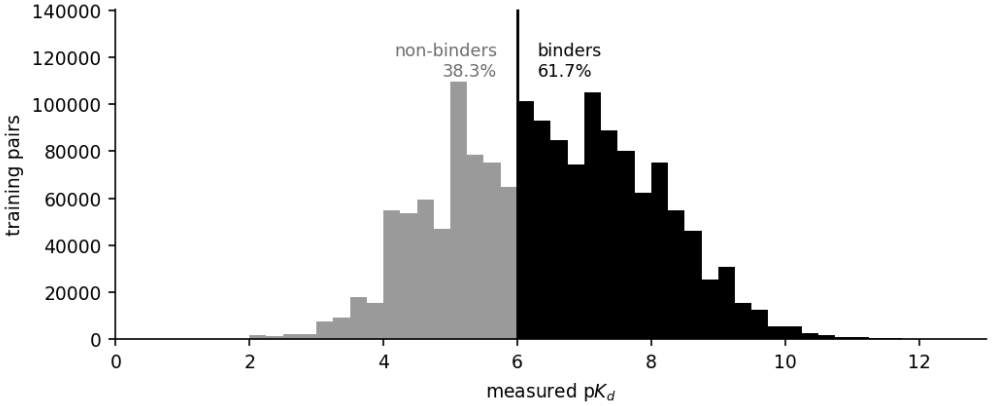
Distribution of the training label across all 1,568,316 collapsed protein–ligand pairs, divided at the pKd 6 binder threshold.

### Structure-free model

For the 1.8 million complexes, we encoded the proteins with ESM2-650M (28), which produces a 1,280-dimensional representation. The ligands were encoded with ChemBERTa-77M-MLM (31), which produces 384 dimensions. We then mean-pooled both over the last hidden state with padding masked out. To these, we added the surviving RDKit named descriptor set, an ECFP4 count fingerprint (48), and amino acid composition with chain length. Table 1 gives the resulting layout.

**Table 1.** Feature layout of the reported model. Block order is load-bearing and must match the order used in training.

| **Block** | **Source** | **Width** |
|---|---|---|
| ChemBERTa-77M-MLM ligand embedding | SMILES | 384 |
| RDKit named descriptors | SMILES | 226 |
| ECFP4 count fingerprint | SMILES | 1,024 |
| ESM2-650M protein embedding | AA sequence | 1,280 |
| Amino acid composition and length | AA sequence | 21 |
| Total Features |  | 2,935 |

### Model training

We fit the model with XGBoost (39) under squared-error loss and a protein-grouped split, in which no protein is shared between the training and test sets. The inner split used for early stopping holds out the same grouping key as the outer split. A random inner split would allow the model to tune against proteins on which it is subsequently tested, which inflates the reported score without improving the model.

We averaged replicate measurements of the same sequence and SMILES pair into a single label before fitting, so that a frequently re-measured pair does not receive proportionally more weight than a singly measured one. We also removed benchmark ligands from the training set before fitting, dropping every training ligand within an ECFP4 Tanimoto similarity of 0.50 of any benchmark ligand, so that the benchmark cannot measure memorization.

The model ships as a regressor alone. We make binder calls by thresholding the predicted pKd, which has the additional advantage that the threshold is selected at inference time and is not fixed during training.

Table 2 reports the held-out performance the model reached during training. This test set is a protein-grouped hold-out drawn from the training corpus, so it measures generalization to unseen proteins within the same data distribution. It is not the external benchmark, and the two should not be compared directly.

**Table 2.** Held-out performance of the reported model during training, on a protein-grouped test split of the training corpus.

| **Quantity** | **Value** |
|---|---|
| Features | 2,935 |
| Fitting pairs | 1,022,678 |
| Validation pairs | 229,846 |
| Held-out test pairs | 315,792 |
| Boosting rounds at best iteration | 1,784 |
| Test *R*<sup>2</sup> | 0.307 |
| Test Pearson *r* | 0.564 |
| Test Spearman *ρ* | 0.536 |
| Test RMSE | 1.270 |
| Test MAE | 1.026 |
| Constant-baseline RMSE | 1.526 |
| Predicted / true spread | 0.508 |

On this split, the model explains 30.7% of the variance in pKd and beats the constant baseline on RMSE at 1.270 against 1.526. Both figures are stronger than the corresponding ones on the external benchmark reported below, which is the expected consequence of a hold-out drawn from the same distribution and the same period as the training data.

### Benchmark construction

To evaluate the model, we assembled a benchmark of 500 protein–ligand pairs across 20 targets from ChEMBL (49, 50). We constructed the set to be difficult in 3 specific respects.

- **Ligand novelty**. Every benchmark ligand falls below an ECFP4 Tanimoto similarity of 0.50 to anything in Savant’s training set. The realized maximum is 0.494.
- **Temporal separation**. Every measurement dates from 2025 or later and is absent from ChEMBL v34 (April 2024), the release most plausibly contained within the training window of the comparison methods.
- **Assay filtering**. We retained *K*_*d*_ and *K*_*i*_ measurements only, at a ChEMBL confidence score of 8 or greater.

This design is a time-split in the sense of Sheridan (51), who showed that random cross-validation is optimistic and that splitting on measurement date estimates prospective performance far more realistically. Combined with the near-neighbor removal, this method implements directly what the memorization literature recommends for benchmarks intended to measure generalization (34, 35).

We deliberately did not require novel proteins. The training corpus covers essentially every well-measured target, so a novel-protein requirement would have left only sparsely characterized targets, for which the ground truth is itself unreliable. All 20 benchmark targets therefore appear in training with other ligands. This benchmark measures the ranking of novel chemistry against an established target, which is the realistic screening scenario, and results obtained from it must not be described as zero-shot with respect to the target.

The temporal filter protects every method equally, because no method trained before 2025 would have seen these measurements. The ligand similarity filter does not transfer between methods, and the direction in which this cuts is worth stating explicitly. We applied the near-neighbor removal to Savant’s training set only. The comparison methods were trained by other groups on other corpora and may retain close analogues of these benchmark ligands. Savant is therefore held to a stricter novelty standard than the methods against which we compare it, and any advantage it shows is measured under that handicap.

We audited the training corpus against the benchmark before every reported run, computing for each benchmark ligand its maximum ECFP4 Tanimoto similarity to the training set, whether its InChIKey skeleton appears in training, and whether the exact pair appears in training. We report a result only when all three checks return zero violations.

### Comparison methods

Table 3 lists the methods compared and the input each requires at inference time.

**Table 3.** The methods compared, and the input each requires at inference time.

| **Method** | **Description** | **Requires** |
|---|---|---|
| **Savant** | This work. Gradient-boosted trees over protein and ligand language-model embeddings, physicochemical descriptors, and a topological fingerprint | sequence and SMILES, CPU |
| **Boltz-2** | Co-folds the protein–ligand complex with a diffusion model, then predicts affinity from the predicted structure | sequence and SMILES, GPU |
| **PRODIGY-LIG** | Physics-based rescoring. A linear model over atomic contact counts in an existing complex | 3-dimensional complex, CPU |
| **PLAPT** | The closest published peer. Protein and chemical language-model embeddings passed to a small neural head, also structure-free | sequence and SMILES, CPU or GPU |

Boltz-2 represents the co-folding approach and is the strongest generally available affinity predictor we could run (19). PRODIGY-LIG represents physics-based rescoring and requires a structure, which we supplied from the same folding workflow used in the structure-based stage (52). PLAPT is the closest structure-free peer (27) and is, at the time of writing, a preprint that has not undergone independent peer review.

### Evaluation and statistical analysis

We decomposed the pooled Pearson correlation into a between-target component, which correlates predicted target means against measured target means, and a within-target component, which centers both quantities on the target mean beforehand. The first corresponds to determining which protein binds more tightly and the second to determining which ligand binds more tightly for a given target. This decomposition is not a standard named analysis in this field. We introduce the decomposition here as a quantitative operationalization of the qualitative warnings that pooled benchmarks reward exploiting target identity (33–36).

We computed Spearman *ρ* within every target carrying at least 10 pairs and averaged across targets. We deliberately do not report enrichment factors. Truchon and Bayly (53) showed that the enrichment factor is insensitive to ranking performance before and after its cutoff. Our benchmark is 88.6% binders, a density at which enrichment saturates near 1.13 and cannot separate a perfect ranker from a mediocre one.

We computed all paired comparisons as cluster-resampled bootstraps, resampling targets instead of individual pairs, so that the resulting interval reflects uncertainty over the units actually sampled. We report Wilcoxon signed-rank and sign tests alongside each bootstrap. All confidence intervals are 95%, and we report a point estimate whose interval spans zero as parity, irrespective of the sign of the estimate. For paired comparisons of within-target correlations, we additionally applied the test reported in Steiger 1980, which accounts for the dependence between two correlations measured against the same values and is therefore more sensitive than comparing two independent intervals.

### Runtime measurement

We measured runtime on CPU, with the protein encoder forced onto the CPU explicitly, because the presence of a visible GPU otherwise alters the measurement even when the GPU is not intended to be used. CPU run time, on an Intel Xeon Gold 6326 @ 2.90GHz, was determined using run statistics from High Performance Cluster (HPC) job manager. We measured 3 quantities separately, because they scale differently: the one-off cost of encoding a protein, the marginal cost of one additional ligand scored against an already-encoded protein, and the total for a realistic mixed workload. We did not take a head-to-head wall-clock measurement of Boltz-2. The runtime comparison reported below is therefore structural, describing the work each method must perform per pair, and is not a measured speedup ratio.

## Results

### Methods do not predict absolute affinity better than a constant

The evaluation set holds 500 pairs across 20 targets and 373 distinct ligand skeletons, with a mean pKd of 7.296 and a standard deviation of 1.274. The realized maximum ECFP4 Tanimoto to anything in training is 0.494. That label spread of 1.274 pKd is the value against which every RMSE below should be judged, because it is precisely the RMSE achieved by predicting the benchmark mean for every pair. Table 4 reports every method against it.

**Table 4.** The four compared methods on the 500-pair benchmark. Between-target *r* corresponds to determining which protein binds more tightly and within-target *r* to determining which ligand binds more tightly for a given target. Mean *ρ* is the within-target Spearman correlation averaged over the 20 targets. Boltz-2 and PRODIGY-LIG returned no prediction for four pairs.

| **Method** | *n* | **Between *r*** | **Within *r*** | **Pooled *r*** | **Mean *ρ*** | **RMSE** |
|---|---|---|---|---|---|---|
| Savant | 500 | 0.100 | **0.436** | 0.227 | **0.278** | 1.353 |
| Boltz-2 | 496 | **0.351** | 0.386 | **0.412** | 0.228 | 1.388 |
| PRODIGY-LIG | 496 | 0.295 | 0.237 | 0.333 | 0.233 | 1.537 |
| PLAPT | 500 | −0.225 | 0.208 | 0.021 | 0.067 | 1.827 |
| Constant baseline | 500 | — | — | — | — | **1.273** |

Two findings follow from this table, and both concern the benchmark itself more than any individual method.

First, no method outperformed the constant baseline on RMSE. Savant’s RMSE of 1.353 is the lowest of any method tested and is lower than Boltz-2’s 1.388, but it remains higher than the 1.273 obtained by predicting 7.296 for every pair. We therefore report no absolute-error claim for any method evaluated on this benchmark, including our own.

This outcome is what the label precision predicts, following the argument of Kramer et al. (38) that experimental uncer-tainty sets an upper limit on achievable performance. Treating the replicate disagreement of 0.81 pKd as the noise standard deviation and the benchmark label spread of 1.274 pKd as the total, the implied signal variance is 1.274^2^ *−* 0.81^2^ = 0.97, so the reliability ratio is 0.97*/*1.62 = 0.60. A model that recovered the true affinity perfectly would therefore reach an *R*^2^ of approximately 0.60 and an RMSE of approximately 0.81 on this benchmark, not 1.0 and 0. The negative *R*^2^ values we observe reflect a benchmark whose dynamic range is narrow relative to its measurement noise, and they are not by themselves evidence that a method is uninformative.

Second, Savant’s pooled correlation of 0.227 trails both Boltz-2 (0.412) and PRODIGY-LIG (0.333). Savant also produces the most compressed predictions of any method tested, at 0.425 of the measured spread. This compression is a consequence of fitting squared-error loss against a label whose own replicate disagreement is 0.81 pKd, which makes a prediction near the conditional mean the loss-minimizing choice. Because rank order is invariant to a change of scale, the compression does not affect any of the ranking metrics reported below. The compression does mean that compounds should be selected by rank, such as the top 1% of a library, and not by an absolute predicted cutoff.

Fig. 2, illustrates that compression pair by pair, against Boltz-2 on the same measurements. We applied regression predicted on measured pKd yields a slope of 0.10 for Savant and 0.31 for Boltz-2. A perfectly calibrated method would give 1.0. Savant’s binned means run from 6.0 to 7.0 across a measured range of 8 log units, so its prediction moves in the right direction but by a fraction of the true distance. Boltz-2 tracks the identity line more closely and disperses its predictions more widely, at the cost of substantially more scatter about the trend. The 2 panels make the trade off explicit: neither method is calibrated, and the one closer to calibrated is not the one that orders compounds better.

**Fig 2.**
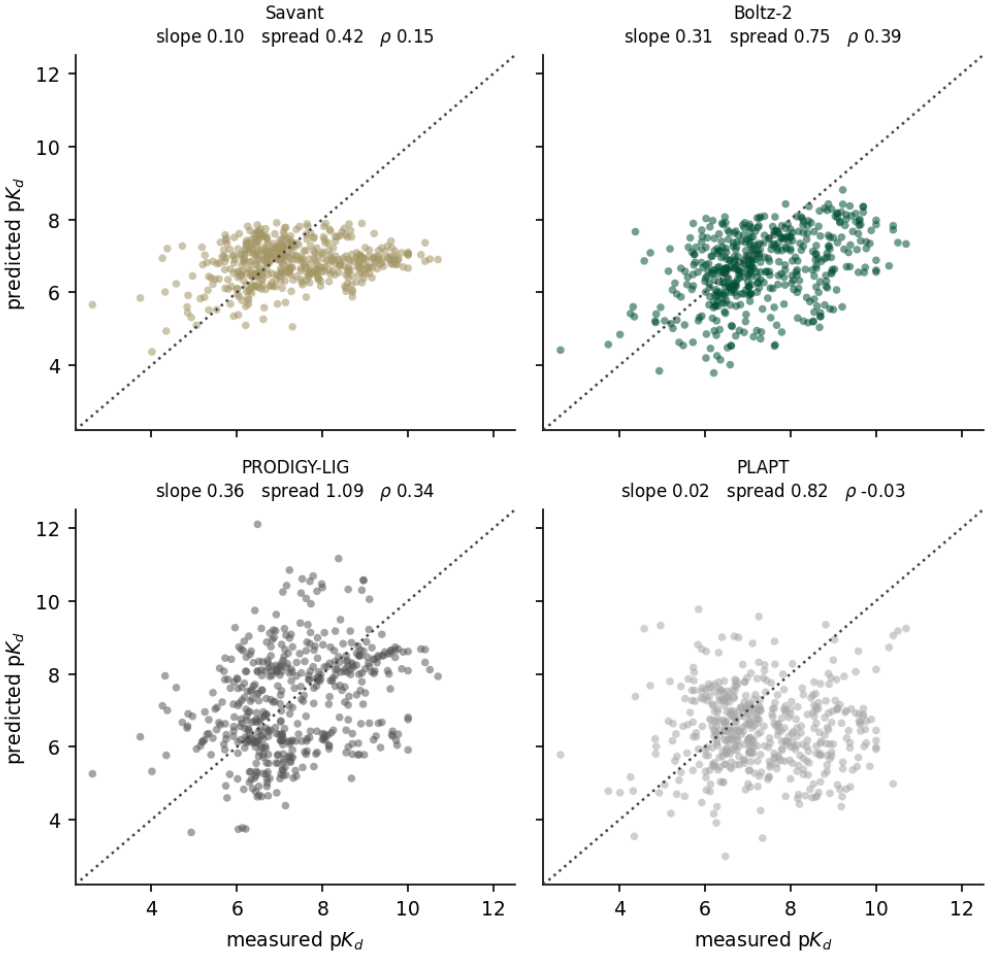
Predicted against measured pKd for Savant and Boltz-2 on the benchmark. The dashed line is the identity. Black markers give the mean prediction within each 1-unit bin of measured affinity, with the standard error of that mean. Slope is the least-squares fit of predicted on measured, spread is the ratio of predicted to measured standard deviation, and *ρ* is the pooled Spearman correlation.

### Decomposition reveals that Savant’s accuracy is with-in-target

To identify the source of Savant’s pooled deficit, we decomposed the pooled correlation into its between-target and within-target components. In Fig. 3, we see the result alongside the calibration figures.

**Fig 3.**
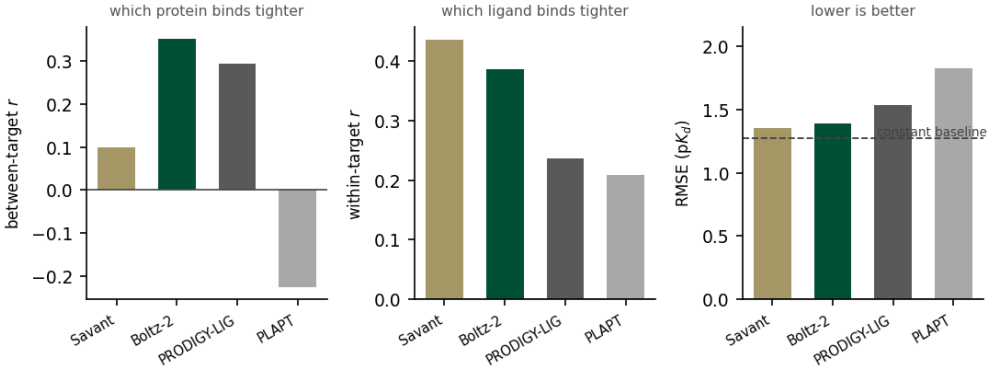
The 4 compared methods on the 3 quantities that matter. Savant achieves the highest within-target correlation and nearly the lowest between-target correlation, whereas Boltz-2 is balanced across the 2. The pooled correlation in Table 4 is a weighted blend of the first 2 panels, and the weighting is a property of the bench-mark. In the third panel every method sits above the constant baseline.

**Fig 4.**
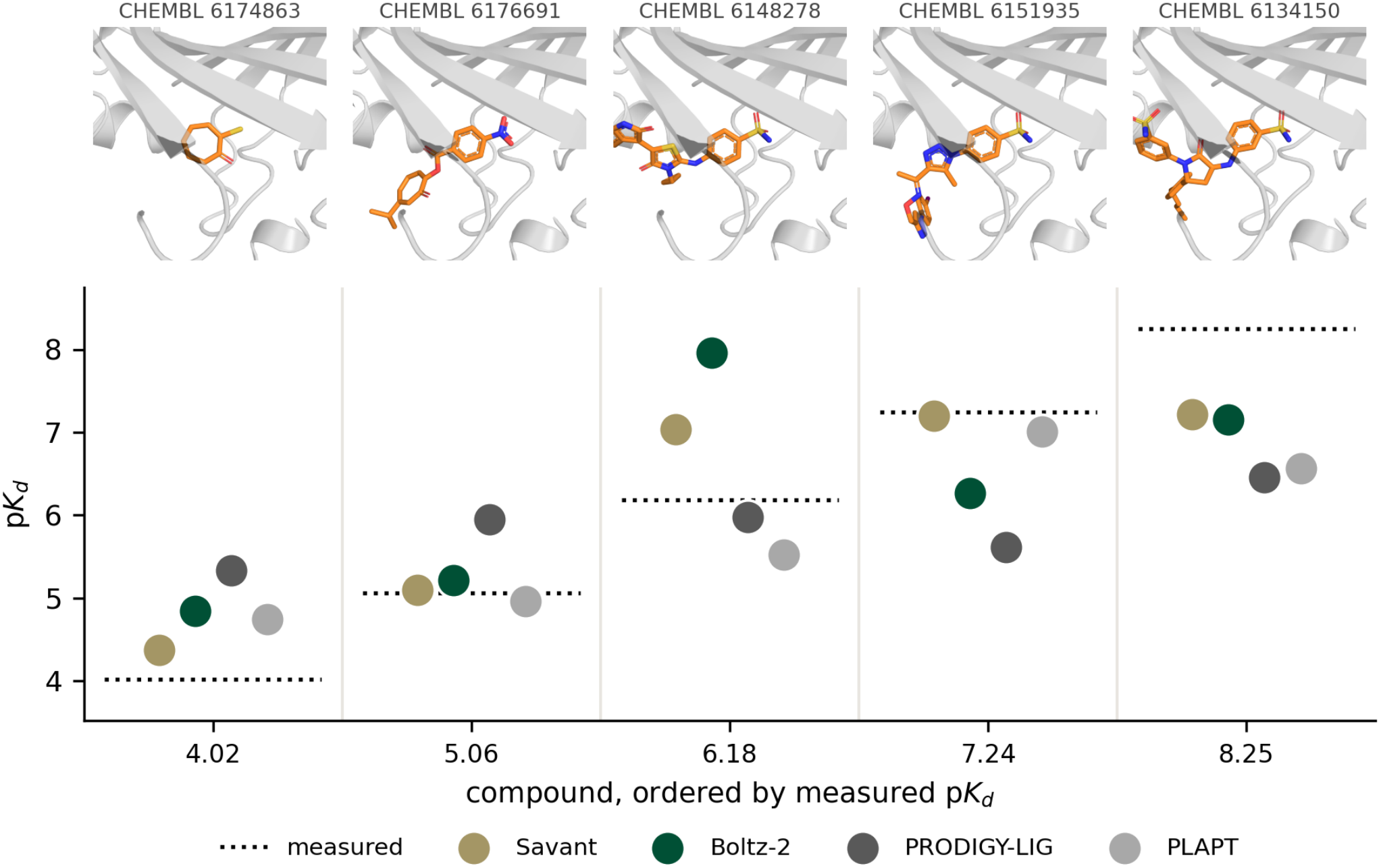
Five carbonic anhydrase 2 complexes, ordered by measured affinity. Top: the Boltz-2 co-folded complex for each ligand with the protein in grey and the ligand in orange. All 5 models are confidently folded, with complex pLDDT between 0.967 and 0.978 (mean 0.972). **Bottom:** true pKd values against the values predicted by Savant and Boltz-2. Savant is closer to the measurement than Boltz-2 for all 5 ligands, with a mean absolute error of 0.46 against 0.97, and its predictions increase with the measurement, whereas Boltz-2 overshoots at pKd 6.18 and crosses the measured line twice.

This is the central result of the evaluation. Savant’s within-target correlation of 0.436 is the highest of any method tested and exceeds Boltz-2’s 0.386. Conversely, Savant’s between-target correlation of 0.100 is close to zero and far below Boltz-2’s 0.351. The pooled deficit is therefore attributable entirely to the between-target component, and on the within-target component Savant leads. These findings indicate that Savant ranks compounds against a given target at least as well as a method that folds every complex on a GPU, and that it does not reliably determine which of 2 proteins binds more tightly.

We tested these differences with Steiger’s paired test on the rows for which both methods returned a prediction. Savant’s advantage over PRODIGY-LIG (*P* = 0.0002) and over PLAPT (*P <* 0.0001) is larger than sampling noise. Its 0.046 advantage over Boltz-2 is not (*P* = 0.24), which is the basis on which we describe the 2 methods as at parity on this axis. We note that the between-target axis of this benchmark is adversarial by construction. Because we selected pairs to span affinity bins within each target, the per-target mean affinity of the benchmark correlates at *r* = *−* 0.17 with the per-target median of the training corpus. A model is therefore penalized on this axis for correctly learning a target’s average level. This property explains part, though not all, of the between-target deficit, and it is a reason to prefer the decomposed figures over the pooled one.

### Savant ranks compounds within a target as well as Boltz-2

To evaluate the screening task directly, we computed Spearman *ρ* within each of the 20 benchmark targets and averaged across them, which is the Mean *ρ* column of Table 4. Savant recorded the highest value of any method tested, at 0.278 against Boltz-2’s 0.228, and a positive correlation on 16 of 20 targets against Boltz-2’s 15.

A cluster bootstrap over targets puts the paired difference against Boltz-2 at +0.050, with a 95% confidence interval from *−* 0.059 to 0.159. The interval spans zero, so we report this as parity: with 20 targets the resolution available does not separate 0.278 from 0.228. The same comparison against PRODIGY-LIG gives +0.005 (*−* 0.144 to 0.176) and against

PLAPT *−*0.161 (*−* 0.305 to *−* 0.019). What the comparison establishes is that a model requiring no structure and no GPU is not measurably worse at the screening task than a model that folds every complex.

### External validation on an independently assembled benchmark

Our custom-built benchmark spans 20 targets, which bounds both its resolution and its impartiality. To test the method against neither constraint, we scored Savant on the leakage-filtered ChEMBL35 set released with PLABench (55, 56), an independent benchmark of 4,862 pairs across 71 targets on which 9 other methods have already been scored by their authors.

PLABench reports a target-weighted mean of per-target correlations rather than a pooled figure. We reproduced all 9 of their published values from their own per-target files before scoring Savant, so a protocol mismatch would appear here as a reproduction difference. Table 5 reports every method on their full dataset, ordered by RMSE.

**Table 5.** The 9 PLABench methods and Savant on their leakage-filtered ChEMBL35 set of 4,862 pairs across 71 targets, under PLABench’s protocol, in which Pearson, Spearman and RMSE are target-weighted means of per-target values. Rows are ordered by RMSE.

| **Method** | **Pearson** | **Spearman** | **RMSE** |
|---|---|---|---|
| FLOWR.ROOT | 0.392 | 0.374 | 0.928 |
| **Savant** | **0.252** | **0.226** | **1.007** |
| MFE | 0.177 | 0.162 | 1.212 |
| BAPred | 0.157 | 0.131 | 1.221 |
| Boltz-2 | 0.321 | 0.302 | 1.248 |
| MixingDTA | 0.173 | 0.162 | 1.272 |
| Haiping | 0.146 | 0.128 | 1.341 |
| FlowDock | 0.188 | 0.172 | 1.369 |
| DeepDTA | 0.153 | 0.151 | 1.394 |
| LLF | 0.212 | 0.209 | 1.462 |

Savant places 2nd of the 10 methods on root mean squared error and 3rd on correlation. Its RMSE of 1.007 is the lowest of any method except FLOWR.ROOT’s (57) 0.928, and well below Boltz-2’s 1.248, both of which require a structure and a GPU for acceleration to determine binding affinity. On correlation, Savant leads every other method that operates without a structure, by 0.040 over LLF and by 0.099 over DeepDTA. Savant reaches that position requiring neither a structure nor a GPU at inference.

The leakage filter behind that set removes test ligands within an ECFP4 Tanimoto of 0.85 of any of 14 training corpora. Savant’s corpus is not among the 14, and our own bench-mark is built on a stricter gate of 0.50, so we audited their ligands against our training set directly. No ligand shares an InChIKey skeleton with anything Savant trained on and none reaches 0.85, the closest being 0.849. Their filter therefore transfers to our corpus. However, 52.1% of their ligands sit at or above 0.50.

Most consequential for the interpretation of our own bench-mark, Savant’s between-target correlation on this set is 0.513, against 0.100 on the 500-pair benchmark reported earlier. PLABench did not select its pairs to span affinity bins within each target, so its between-target axis is not adversarial in the way ours is. The near-zero figure we report on our own benchmark is therefore substantially a property of that bench-mark’s construction, and the method ranks targets against one another considerably better than our benchmark alone would suggest.

### The cost of screening is dominated by encoding the protein once

The result here is the cost structure, not the absolute timing. The expensive step is encoding the protein, and it is performed once per target. Each additional ligand costs 0.018s, because the marginal path consists of descrip-tor computation and tree inference, neither of which benefits substantially from a GPU. Screening 1 target against 100,000 compounds is therefore projected at approximately half an hour on a CPU. Table 6 reports measured CPU timings.

**Table 6.** Measured CPU timings, with the protein encoder forced onto CPU.

| **Quantity** | **CPU** |
|---|---|
| Protein encoding, per protein (one-off) | 1.691s |
| Per additional ligand against an encoded protein | 0.018s |
| Cold single pair, end to end | 1.710s |
| 500 pairs across 20 targets (the full benchmark) | 42.9s |
| 1 target against 100,000 ligands (projected) | 0.505hr |

This results constitutes a structural advantage over co-folding methods rather than a constant-factor one. A method that folds the complex must repeat its expensive step for every ligand, so its cost is linear in library size with a large constant. In contrast, Savant’s cost is linear with a small constant plus a fixed offset. We did not take a head-to-head wall-clock measurement against Boltz-2, and we therefore do not report a speedup ratio.

## Discussion

In this study, we built a protein–ligand affinity predictor for drug screening. We then evaluated the resulting structure-free model on a temporally separated, ligand-novel bench-mark against a highly popular co-folding method, a physics-based rescoring function, and the closest structure-free peer. Our central finding is that the ability to predict affinity decomposes into 2 separable abilities that are not equally difficult, and that a method can lead decisively on 1 while trailing on the other.

### The metric selects the winner

The ordering of methods by pooled correlation is close to the inverse of the ordering by within-target correlation. Boltz-2 leads on the pooled figure at 0.412 against Savant’s 0.227, and Savant leads on the within-target figure at 0.436 against Boltz-2’s 0.386. Both numbers are computed on the same 500 pairs. The difference is attributable to how each metric weights the between-target and within-target components, and that weighting is a property of the benchmark rather than of any method.

That the choice of metric can reorder methods is not itself new. Truchon and Bayly (53) made the argument for virtual-screening classification, showing that the enrichment factor and the area under the ROC curve rank methods differently because they weight early recognition differently. The bias literature makes the complementary point that pooled bench-marks can be satisfied by exploiting dataset structure rather than by learning interactions (33–35), and Volkov et al. (36) showed for affinity regression specifically that explicit interaction descriptions add nothing over ligand-only descriptors, with memorization dominating true learning. In this work, we add a quantitative decomposition for the regression setting, and a demonstration that on a contamination-controlled benchmark the inversion is large enough to reverse the ranking of a GPU co-folding method and a CPU tree ensemble. We argue that reporting a single pooled correlation for a protein–ligand affinity method is insufficient, because the pooled figure blends an ability that screening requires with an ability that screening never exercises. A screening campaign fixes the protein. Under that condition the between-target component is not merely less important; it is unobservable, and a method’s score on it carries no information about how it will perform.

### The utility of structure-free

Two independent lines of published evidence support the same conclusion. Volkov et al. (36) found that explicitly describing protein–ligand non-covalent interactions gave no advantage over ligand or protein descriptors alone, which is a strong external statement that the marginal value of structural detail is small. Karelina et al. (21) found that predicted protein structures do not yet support ligand-pose prediction at the accuracy of experimentally determined structures. This result means that the structures available at scale are not the structures suitable for the method. Incorporating structures into the pipeline increases run time and computational resources, thus limiting accessibility for any drug screening lab.

### Ranking is the quantity a screening decision consumes

A screening decision consumes an ordering. When a chemist selects the top 1% of a library for synthesis, that choice is unchanged by any monotone transformation of the predicted values. What matters is how closely the predicted order matches the measured one. *R*^2^ and RMSE answer a different question, namely how much of the variance in the measured values a method reproduces on the original scale. The distinction between ranking quality and calibration is well established in virtual-screening evaluation (53).

That distinction is what reconciles Savant’s leading within-target correlation with the RMSE results shown in Table 4. Measurement noise caps the achievable *R*^2^ on this bench-mark near 0.60 for any method whatsoever, and every method we tested fell below the constant baseline on RMSE, so the calibration figures characterize the benchmark and its label at least as much as they characterize the models. The ranking figures are the ones a campaign acts on, and on those Savant leads: a within-target correlation of 0.436 against Boltz-2’s 0.386, and a positive rank correlation on 16 of the 20 bench-mark targets.

### Where the method applies

Savant is built for the case in which the protein is fixed and many candidate ligands must be ordered, which covers hit triage, library prioritization, and the iterative rounds of early lead optimization. This is the common situation in practice, because a project commits to a target long before it commits to a chemical series, and most decisions that follow concern which molecules to make next. The cost structure has the same shape as the task. Encoding the protein costs 1.691s and happens once per target, after which each additional ligand costs 0.018s. Thus, a library of 100,000 compounds can be scored in approximately half an hour on a CPU. A campaign that rescores its library after each synthetic round pays the protein cost once and the ligand cost repeatedly, which is the regime in which the advantage over a method that folds every complex compounds with every round.

Savant requires no structural input of any kind. Savant extends to targets for which no experimental structure exists and for which a predicted structure would not yet support quantitative use (21). Running on a CPU, Savant can be em-bedded directly in a generative design loop, where candidate molecules are proposed and scored in the same process with-out contending for allocation of an accelerator.

The benchmark measured this scenario under stringent conditions. Every ligand fell below an ECFP4 Tanimoto of 0.50 to anything in training. Every measurement postdated the training window of the methods compared, thus the reported performance is on genuinely novel chemistry against established targets. That is a common problem in a screening campaign and represents the setting in which published performance most often degrades (34, 51). Under those conditions, Savant matched a method that folds every complex on a GPU, and it achieved a positive within-target rank correlation on 16 of 20 targets.

Two scope conditions follow from the same decomposition and the external validation qualifies the first of them. Ranking candidate protein targets for a fixed ligand is a different question and Savant ranks targets less well than it ranks ligands. The size of that gap depends on the benchmark: the near-zero 0.100 we record on our own set rises to 0.513 on PLABench. PLABench’s between-target axis is not adversarially constructed, so the method is weaker on this axis rather than blind to it. Neither condition bears on the screening case in which the protein is fixed and the ordering is what the decision consumes.

### Relation to other structure-free methods

PLAPT (27), which is similar to Savant, also predicts affinity from sequence and SMILES using protein language-model embeddings. On this benchmark, PLAPT recorded a pooled correlation of 0.021 and a between-target correlation of *−* 0.225. We do not interpret this as evidence that structure-free prediction is intrinsically weak, since Savant is also structure-free and records a within-target correlation of 0.436 on the same pairs. The comparison suggests that feature construction, machine learning algorithm selection, and the choice of training objective matter more than the presence or absence of a structure. This conclusion is shared by Volkov et al. (36). Published performance in this literature is known to depend heavily on how the evaluation is split. Random splits produce systematically optimistic estimates (34, 51).

We evaluate every method here on the same temporally separated, ligand-novel pairs, which is a more stringent condition than the benchmarks on which these methods were originally tested.

### Evaluation design

We regard the benchmark design as a contribution independent of the model. Requiring measurements published after the training window of every method implements a time split in Sheridan’s (51) sense, in which he showed estimates prospective performance far more realistically than random cross-validation. Requiring every bench-mark ligand to fall below an ECFP4 Tanimoto of 0.50 to any-thing in training addresses the memorization failure mode that Wallach and Heifets (34) documented for ligand-based benchmarks. Holding out the inner validation fold on the same grouping key as the outer split extends the same principle to model selection such that early stopping is not tuned on proteins on which the model will be tested.

## Limitations

The breadth of the data on which we could assemble and train is limiting. The training corpus for Savant is far less biologically diverse than its size suggests. The corpus holds 1,836,729 measurements over 22,974 proteins and 991,876 ligands. However, the protein coverage is heavily concentrated. 224 proteins account for half of all measurements, 1,576 account for 90%, the median protein carries two measurements, and 77.4% of proteins carry fewer than 10. Most of what the model learns about proteins comes from a narrow slice of well-studied targets. Assembling a broader and more balanced corpus and evaluating over more targets will require gathering and curating measurements on a scale we could not reach within this project timeline. Performance on some sparsely measured protein families is consequently not established. We shall seek resources to do these measurements in the future.

## Conclusions

We set out to determine whether binding affinity could be predicted for ligand screening without using or predicting a 3-dimensional protein structure and without a GPU. These two requirements are what make use of existing accurate methods difficult. We find that our goal can be achieved provided the prediction is used as an ordering.

Savant takes an amino acid sequence and a SMILES string and returns a predicted pKd. Savant ranks ligands against a fixed protein as well as other tools that needs folding of every complex on a graphics processing unit (i.e. Boltz-2), at a within-target correlation of 0.436 against 0.386, a difference the paired test places within sampling noise. In a comparison of Savant against the physics-based and structure-free comparators (e.g., Boltz-2, PRODIGY-LIG, PLAPT) the margin is larger than noise. With Savant, encoding a protein costs 1.691s and happens once per target, after which each additional ligand costs 0.018s. It follows that a library of 100,000 compounds can be ranked in half an hour on a central processing unit. Because no 3-dimensional protein structure is required at any point with Savant, the method applies to targets for which these data are lacking. Compounds can be selected from Savant’s output by rank, such as the top 1% of a library. The predicted values should not be read as calibrated dissociation constants. What Savant offers is a prediction accurate enough for the ranking decision needed in a screening campaign, but delivered at a reduced computational cost over other methods.

In the end, this contribution includes a screening model that runs on commodity hardware and a contamination-controlled benchmark on which it and 3 published methods are scored under a single implementation.

## Code and Data Availability

All dataset generation code, model training logic, machine learning models, and evaluations are available at https://github.com/sayalg/Savant.

## Author Contributions

SGM conceptualized the study, designed and built Savant and the benchmark, performed the data analysis, and drafted the manuscript. CTF contributed to the organization of the study, the development of initial machine learning approach and evaluation logic, and writing the manuscript. DJ contributed to the writing and editing of the manuscript.

## Acknowledgments

We acknowledge the following entities at the University of North Carolina at Charlotte: Academic Affairs, The Division of Research (Ignite Award), The Center for Computational Intelligence to Predict Health and Environmental Risks (CI-PHER), The Department of Bioinformatics and Genomics, The College of Computing and Informatics, and the University Research Computing group.

This work was built on public data, models, and code released by others. The training corpus was compiled by Jens Glaser and hosted on Hugging Face. We thank Meta AWe Research for ESM2, the DeepChem project for ChemBERTa, Google DeepMind for AlphaFold3, the Bonvin laboratory for HAD-DOCK3 and PRODIGY-LIG, and the authors of Boltz-2 and PLAPT. The external validation set and the predictions of the nine methods scored against it were released by LyuweWe Wang and Jianlin Cheng.

## Bibliography

1. Olivier J. Wouters, Martin McKee, and Jeroen Luyten. Estimated research and development investment needed to bring a new medicine to market, 2009–2018. JAMA, 323(9):844–853, 2020. doi: 10.1001/jama.2020.1166.

2. Joseph A. DiMasi, Henry G. Grabowski, and Ronald W. Hansen. Innovation in the pharmaceutical industry: New estimates of R&D costs. Journal of Health Economics, 47:20–33, 2016. doi: 10.1016/j.jhealeco.2016.01.012.

3. Jack W. Scannell, Alex Blanckley, Helen Boldon, and Brian Warrington. Diagnosing the decline in pharmaceutical R&D efficiency. Nature Reviews Drug Discovery, 11(3):191–200, 2012. doi: 10.1038/nrd3681.

4. Steven M. Paul, Daniel S. Mytelka, Christopher T. Dunwiddie, Charles C. Persinger, Bernard H. Munos, Stacy R. Lindborg, and Aaron L. Schacht. How to improve R&D productivity: The pharmaceutical industry’s grand challenge. Nature Reviews Drug Discovery, 9(3):203–214, 2010. doi: 10.1038/nrd3078.

5. Jiankun Lyu, Sheng Wang, Trent E. Balius, Isha Singh, Anat Levit, Yurii S. Moroz, Matthew J. O’Meara, Tao Che, Enkhjargal Algaa, Kateryna Tolmachova, Andrey A. Tolmachev, Brian K. Shoichet, Bryan L. Roth, and John J. Irwin. Ultra-large library docking for discovering new chemotypes. Nature, 566(7743):224–229, 2019. doi: 10.1038/s41586-019-0917-9.

6. Christoph Gorgulla, Andras Boeszoermenyi, Zi-Fu Wang, Patrick D. Fischer, Paul W. Coote, Krishna M. Padmanabha Das, Yehor S. Malets, Dmytro S. Radchenko, Yurii S. Moroz, David A. Scott, Konstantin Fackeldey, Moritz Hoffmann, Iryna Iavniuk, Gerhard Wagner, and Haribabu Arthanari. An open-source drug discovery platform enables ultra-large virtual screens. Nature, 580(7805):663–668, 2020. doi: 10.1038/s41586-020-2117-z.

7. Benjamin I. Tingle, Khanh G. Tang, Mar Castanon, John J. Gutierrez, Munkhzul Khurelbaatar, Chinzorig Dandarchuluun, Yurii S. Moroz, and John J. Irwin. ZINC-22 — a free multi-billion-scale database of tangible compounds for ligand discovery. Journal of Chemical Information and Modeling, 63(4):1166–1176, 2023. doi: 10.1021/acs.jcim.2c01253.

8. Marcus Olivecrona, Thomas Blaschke, Ola Engkvist, and Hongming Chen. Molecular denovo design through deep reinforcement learning. Journal of Cheminformatics, 9:48, 2017. doi: 10.1186/s13321-017-0235-x. Article number 48 — verify if precision critical.

9. Thomas Blaschke, Josep Arús-Pous, Hongming Chen, Christian Margreitter, Christian Tyrchan, Ola Engkvist, Kostas Papadopoulos, and Atanas Patronov. REINVENT 2.0: An AI tool for de novo drug design. Journal of Chemical Information and Modeling, 60(12):5918–5922, 2020. doi: 10.1021/acs.jcim.0c00915.

10. Anastasiia V. Sadybekov and Vsevolod Katritch. Computational approaches streamlining drug discovery. Nature, 616(7958):673–685, 2023. doi: 10.1038/s41586-023-05905-z.

11. Michael K. Gilson and Huan-Xiang Zhou. Calculation of protein–ligand binding affinities. Annual Review of Biophysics and Biomolecular Structure, 36:21–42, 2007. doi: 10.1146/annurev.biophys.36.040306.132550.

12. Andrew L. Hopkins, Colin R. Groom, and Alexander Alex. Ligand efficiency: A useful metric for lead selection. Drug Discovery Today, 9(10):430–431, 2004. doi: 10.1016/S1359-6446(04)03069-7.

13. Gregory L. Warren, C. Webster Andrews, Anna-Maria Capelli, Brian Clarke, Judith LaLonde, Millard H. Lambert, Mika Lindvall, Neysa Nevins, Simon F. Semus, Stefan Senger, Giovanna Tedesco, Ian D. Wall, James M. Woolven, Catherine E. Peishoff, and Martha S. Head. A critical assessment of docking programs and scoring functions. Journal of Medicinal Chemistry, 49(20):5912–5931, 2006. doi: 10.1021/jm050362n.

14. Minyi Su, Qifan Yang, Yu Du, Guoqin Feng, Zhihai Liu, Yan Li, and Renxiao Wang. Comparative assessment of scoring functions: The CASF-2016 update. Journal of Chemical Information and Modeling, 59(2):895–913, 2019. doi: 10.1021/acs.jcim.8b00545.

15. Lingle Wang, Yujie Wu, Yuqing Deng, Byungchan Kim, Levi Pierce, Goran Krilov, Dmitry Lupyan, Shaughnessy Robinson, Markus K. Dahlgren, Jeremy Greenwood, et al. Accurate and reliable prediction of relative ligand binding potency in prospective drug discovery by way of a modern free-energy calculation protocol and force field. Journal of the American Chemical Society, 137(7):2695–2703, 2015. doi: 10.1021/ja512751q.

16. Christina E. M. Schindler, Hannah Baumann, Andreas Blum, Dietrich Boöse, Hans-Peter Buchstaller, et al. Large-scale assessment of binding free energy calculations in active drug discovery projects. Journal of Chemical Information and Modeling, 60(11):5457–5474, 2020. doi: 10.1021/acs.jcim.0c00900. Verify full author list and pages.

17. Josh Abramson, Jonas Adler, Jack Dunger, Richard Evans, Tim Green, Alexander Pritzel, Olaf Ronneberger, Lindsay Willmore, Andrew J. Ballard, Joshua Bambrick, et al. Accurate structure prediction of biomolecular interactions with AlphaFold 3. Nature, 630(8016):493–500, 2024. doi: 10.1038/s41586-024-07487-w.

18. Jeremy Wohlwend, Gabriele Corso, Saro Passaro, Mateo Reveiz, Ken Leidal, Wojtek Swiderski, Tally Portnoi, Itamar Chinn, Jacob Silterra, Tommi Jaakkola, and Regina Barzilay. Boltz-1: Democratizing biomolecular interaction modeling. bioRxiv, 2024. doi: 10.1101/2024.11.19.624167. Preprint.

19. Saro Passaro, Gabriele Corso, Jeremy Wohlwend, Mateo Reveiz, Stephan Thaler, Vignesh Ram Somnath, Noah Getz, Tally Portnoi, Julien Roy, Hannes Stärk, David KwabiAddo, Dominique Beaini, Tommi Jaakkola, and Regina Barzilay. Boltz-2: Towards accurate and efficient binding affinity prediction. bioRxiv, 2025. doi: 10.1101/2025.06.14.659707. Preprint.

20. Gabriele Corso, Hannes Stärk, Bowen Jing, Regina Barzilay, and Tommi Jaakkola. DiffDock: Diffusion steps, twists, and turns for molecular docking. In International Conference on Learning Representations (ICLR), 2023. arXiv:2210.01776; no journal DOI.

21. Masha Karelina, Joseph J. Noh, and Ron O. Dror. How accurately can one predict drug binding modes using AlphaFold models? eLife, 12:RP89386, 2023. doi: 10.7554/eLife.89386.

22. Hakime Öztürk, Arzucan Özgür, and Elif Ozkirimli. DeepDTA: Deep drug–target binding affinity prediction. Bioinformatics, 34(17):i821–i829, 2018. doi: 10.1093/bioinformatics/bty593.

23. Thin Nguyen, Hang Le, Thomas P. Quinn, Tri Nguyen, Thuc Duy Le, and Svetha Venkatesh. GraphDTA: Predicting drug–target binding affinity with graph neural networks. Bioinformatics, 37(8):1140–1147, 2021. doi: 10.1093/bioinformatics/btaa921.

24. Kexin Huang, Cao Xiao, Lucas M. Glass, and Jimeng Sun. MolTrans: Molecular interaction transformer for drug–target interaction prediction. Bioinformatics, 37(6):830–836, 2021. doi:10.1093/bioinformatics/btaa880.

25. Rohit Singh, Samuel Sledzieski, Bryan Bryson, Lenore Cowen, and Bonnie Berger. Contrastive learning in protein language space predicts interactions between drugs and protein targets. Proceedings of the National Academy of Sciences, 120(24):e2220778120, 2023. doi: 10.1073/pnas.2220778120.

26. Peizhen Bai, Filip Miljković, Bino John, and Haiping Lu. Interpretable bilinear attention network with domain adaptation improves drug–target prediction. Nature Machine Intelligence, 5(2):126–136, 2023. doi: 10.1038/s42256-022-00605-1.

27. Tyler Rose, Nicolò Monti Navvye Anand, and Tianyu Shen. PLAPT: Protein-ligand binding affinity prediction using pretrained transformers. bioRxiv, 2024. doi: 10.1101/2024.02.08.575577. Preprint — not peer-reviewed.

28. Zeming Lin, Halil Akin, Roshan Rao, Brian Hie, Zhongkai Zhu, Wenting Lu, Nikita Smetanin, Robert Verkuil, Ori Kabeli, Yaniv Shmueli, Allan dos Santos Costa, Maryam Fazel-Zarandi, Tom Sercu, Salvatore Candido, and Alexander Rives. Evolutionary-scale prediction of atomic-level protein structure with a language model. Science, 379(6637):1123–1130, 2023. doi: 10.1126/science.ade2574.

29. EvolutionaryScale Team. ESM cambrian (ESM-C): Scaling protein language models. EvolutionaryScale; open weights and code, 2024. No peer-reviewed paper at the 2024 release; Zenodo software DOI cited for reproducibility.

30. Seyone Chithrananda, Gabriel Grand, and Bharath Ramsundar. ChemBERTa: Largescale self-supervised pretraining for molecular property prediction. arXiv preprint arXiv:2010.09885, 2020. doi: 10.48550/arXiv.2010.09885.

31. Walid Ahmad, Elana Simon, Seyone Chithrananda, Gabriel Grand, and Bharath Ramsundar. ChemBERTa-2: Towards chemical foundation models. arXiv preprint arXiv:2209.01712, 2022. doi: 10.48550/arXiv.2209.01712.

32. Kexin Huang, Tianfan Fu, Wenhao Gao, Yue Zhao, Yusuf Roohani, Jure Leskovec, Connor W. Coley, Cao Xiao, Jimeng Sun, and Marinka Zitnik. Therapeutics data commons: Machine learning datasets and tasks for drug discovery and development. In Advances in Neural Information Processing Systems (NeurIPS), Datasets and Benchmarks Track, 2021. arXiv:2102.09548; no journal DOI.

33. Lieyang Chen, Anthony Cruz, Steven Ramsey, Callum J. Dickson, Jose S. Duca, Viktor Hornak, David R. Koes, and Tom Kurtzman. Hidden bias in the DUD-E dataset leads to misleading performance of deep learning in structure-based virtual screening. PLoS ONE, 14(8):e0220113, 2019. doi: 10.1371/journal.pone.0220113.

34. Izhar Wallach and Abraham Heifets. Most ligand-based classification benchmarks reward memorization rather than generalization. Journal of Chemical Information and Modeling, 58 (5):916–932, 2018. doi: 10.1021/acs.jcim.7b00403.

35. Jochen Sieg, Florian Flachsenberg, and Matthias Rarey. In need of bias control: Evaluating chemical data for machine learning in structure-based virtual screening. Journal of Chemical Information and Modeling, 59(3):947–961, 2019. doi: 10.1021/acs.jcim.8b00712.

36. Mikhail Volkov, Joseph-André Turk Nicolas Drizard, Nicolas Martin, Brice Hoffmann, Yann Gaston-Mathé, and Didier Rognan. On the frustration to predict binding affinities from protein–ligand structures with deep neural networks. Journal of Medicinal Chemistry, 65 (11):7946–7958, 2022. doi: 10.1021/acs.jmedchem.2c00487.

37. Andrea Mastropietro, Giuseppe Pasculli, and Jürgen Bajorath.Learning characteristics of graph neural networks predicting protein–ligand affinities. Nature Machine Intelligence, 5: 1427–1436, 2023. doi: 10.1038/s42256-023-00756-9.

38. Christian Kramer, Tuomo Kalliokoski, Peter Gedeck, and Anna Vulpetti. The experimental uncertainty of heterogeneous public Ki data. Journal of Medicinal Chemistry, 55(11):5165–5173, 2012. doi: 10.1021/jm300131x.

39. Tianqi Chen and Carlos Guestrin. XGBoost: A scalable tree boosting system. In Proceedings of the 22nd ACM SIGKDD International Conference on Knowledge Discovery and Data Mining (KDD ‘16), pages 785–794. ACM, 2016. doi: 10.1145/2939672.2939785.

40. Léo Grinsztajn, Edouard Oyallon, and Gaël Varoquaux. Why do tree-based models still outperform deep learning on typical tabular data? In Advances in Neural Information Processing Systems (NeurIPS), Datasets and Benchmarks Track, volume 35, pages 507–520, 2022. arXiv:2207.08815.

41. Ravid Shwartz-Ziv and Amitai Armon. Tabular data: Deep learning is not all you need. Information Fusion, 81:84–90, 2022. doi: 10.1016/j.inffus.2021.11.011. Verify volume/pages.

42. Jens Glaser et al. jglaser/binding_affinity: Aggregated protein sequence–ligand SMILES binding affinity dataset. Hugging Face Datasets, https://huggingface.co/datasets/jglaser/binding_affinity, 3 2022. No dedicated dataset DOI located; aggregates BindingDB, PDBbind-CN, BioLiP, Binding MOAD. Closest companion paper:Blanchard et al., Int. J. High Perform. Comput. Appl. 2022, 36(5–6):587–602, doi:10.1177/10943420221121804. Confirm intended citation.

43. Michael K. Gilson, Tiqing Liu, Michael Baitaluk, George Nicola, Linda Hwang, and Jenny Chong. BindingDB in 2015: A public database for medicinal chemistry, computational chemistry and systems pharmacology. Nucleic Acids Research, 44(D1):D1045–D1053, 2016. doi: 10.1093/nar/gkv1072.

44. Renxiao Wang, Xueliang Fang, Yipin Lu, and Shaomeng Wang. The PDBbind database: Collection of binding affinities for protein–ligand complexes with known three-dimensional structures. Journal of Medicinal Chemistry, 47(12):2977–2980, 2004. doi: 10.1021/jm030580l.

45. Zhihai Liu, Yan Li, Li Han, Jie Li, Jie Liu, Zhixiong Zhao, Wei Nie, Yuchen Liu, and Renxiao Wang. PDB-wide collection of binding data: Current status of the PDBbind database. Bioinformatics, 31(3):405–412, 2015. doi: 10.1093/bioinformatics/btu626.

46. Jianyi Yang, Ambrish Roy, and Yang Zhang. BioLiP: A semi-manually curated database for biologically relevant ligand–protein interactions. Nucleic Acids Research, 41(D1):D1096–D1103, 2013. doi: 10.1093/nar/gks966.

47. Liegi Hu, Mark L. Benson, Richard D. Smith, Michael G. Lerner, and Heather A. Carlson. Binding MOAD (mother of all databases). Proteins: Structure, Function, and Bioinformatics, 60(3):333–340, 2005. doi: 10.1002/prot.20512.

48. David Rogers and Mathew Hahn. Extended-connectivity fingerprints. Journal of Chemical Information and Modeling, 50(5):742–754, 2010. doi: 10.1021/ci100050t.

49. Anna Gaulton, Louisa J. Bellis, A. Patricia Bento, Jon Chambers, Mark Davies, Anne Hersey, Yvonne Light, Shaun McGlinchey, David Michalovich, Bissan Al-Lazikani, and John P. Overington. ChEMBL: A large-scale bioactivity database for drug discovery. Nucleic Acids Research, 40(D1):D1100–D1107, 2012. doi: 10.1093/nar/gkr777.

50. Barbara Zdrazil, Eloy Felix, Fiona Hunter, Emma J. Manners, James Blackshaw, Sybilla Corbett, Marleen de Veij, Harris Ioannidis, David Mendez Lopez, Juan F. Mosquera, Maria Paula Magarinos, Nicolas Bosc, Ricardo Arcila, Tevfik Kizilören, Anna Gaulton, A. Patrícia Bento, Melissa F. Adasme, Peter Monecke, Gregory A. Landrum, and Andrew R. Leach. The ChEMBL database in 2023: A drug discovery platform spanning multiple bioactivity data types and time periods. Nucleic Acids Research, 52(D1):D1180–D1192, 2024. doi: 10.1093/nar/gkad1004.

51. Robert P. Sheridan. Time-split cross-validation as a method for estimating the goodness of prospective prediction. Journal of Chemical Information and Modeling, 53(4):783–790, 2013. doi: 10.1021/ci400084k.

52. Anna Vangone, Joerg Schaarschmidt, Panagiotis Koukos, Cunliang Geng, Nevia Citro, Mikael E. Trellet, Li C. Xue, and Alexandre M. J. J. Bonvin. Large-scale prediction of binding affinity in protein–small ligand complexes: The PRODIGY-LIG web server. Bioinformatics, 35(9):1585–1587, 2019. doi: 10.1093/bioinformatics/bty816.

53. Jean-François Truchon and Christopher I. Bayly. Evaluating virtual screening methods: Good and bad metrics for the “early recognition” problem. Journal of Chemical Information and Modeling, 47(2):488–508, 2007. doi: 10.1021/ci600426e.

54. James H. Steiger. Tests for comparing elements of a correlation matrix. Psychological Bulletin, 87(2):245–251, 1980. doi: 10.1037/0033-2909.87.2.245.

55. Lyuwei Wang and Jianlin Cheng. PLABench: Data for leakage-controlled benchmarking reveals generalization limits of deep learning for protein–ligand binding affinity prediction. Zenodo, 2026.

56. Lyuwei Wang and Jianlin Cheng. PLABench: A leakage-controlled benchmark for protein–ligand binding affinity prediction. Zenodo, version 1.0.1, 2026.

57. Julian Cremer, Tuan Le, Mohammad M. Ghahremanpour, Emilia Sługocka, Filipe Menezes, and Djork-Arné Clevert. Flowr.root – a flow matching-based foundation model for joint multipurpose structure-aware 3d ligand generation and affinity prediction. Nature Communications, 17(15883), july 2026. doi: 10.1038/s41467-026-74130-9.

